# Heterologous expression and purification of glutamate decarboxylase-1 from the model plant *Arabidopsis thaliana*: characterization of the enzyme’s *in vitro* truncation by thiol endopeptidase activity

**DOI:** 10.1101/2024.04.11.589046

**Authors:** Brittany S. Menard, Kirsten H. Benidickson, Lee Marie Raytek, Wayne A. Snedden, William C. Plaxton

**Affiliations:** Dept. of Biology, Queen’s Univ., Kingston, Ontario, Canada, K7L 3N6; Dept. of Plant Sciences, McGill Univ., Ste-Anne-de-Bellevue, Quebec, Canada, H9X 3V9

**Keywords:** *Arabidopsis thaliana* (Mouse-ear Cress); calcium signaling, calmodulin, γ-aminobutyrate (GABA) shunt, glutamate decarboxylase, proteolysis

## Abstract

Plant glutamate decarboxylase (GAD) is a Ca^2+^-calmodulin activated cytosolic enzyme that produces γ-aminobutyrate (GABA) as the first committed step of the GABA shunt. This pathway circumvents the 2-oxoglutarate to succinate reactions of the mitochondrial tricarboxylic acid cycle. Our prior research established that *in vivo* phosphorylation of the root-specific AtGAD1 isozyme (AT5G17330) occurs at multiple N-terminal serine residues, following Pi resupply to Pi-starved cell cultures of the model plant *Arabidopsis thaliana*. The aim of the current investigation was to purify recombinant AtGAD1 following its expression in *Escherichia coli* to facilitate studies of the impact of site-specific phosphorylation on its kinetic properties. However, *in vitro* proteolytic truncation of a 5 kDa polypeptide from the C-terminus of 59 kDa AtGAD1 subunits occurred during its purification. Immunoblotting demonstrated that most protease inhibitors or cocktails that we tested were ineffective in suppressing partial AtGAD1 proteolysis during incubation of clarified extracts at 23 °C. Although the thiol modifiers N-ethylmaleimide or 2,2-dipyridyl disulfide negated AtGAD1 proteolysis, they also abolished its GAD activity. This indicates that an essential -SH group is needed for catalytic activity, and that AtGAD1 is susceptible to partial degradation either by an *E. coli* cysteine endopeptidase, or possibly via autoproteolytic activity. The inclusion of exogenous Ca^2+^/calmodulin in extraction and chromatography buffers facilitated the purification of non-proteolyzed AtGAD1 to a specific activity of 27 (µmol GABA produced/mg) at optimal pH 5.8, while exhibiting an approximate 3-fold activation by Ca^2+^/CaM at pH 7.3. By contrast, the purified partially proteolyzed His_6_-AtGAD1 was >40% less active at both pH values, and only activated 2-fold by Ca^2+^/CaM at pH 7.3. These results emphasize the need to diagnose and prevent unwanted proteolysis before conducting kinetic studies of purified regulatory enzymes.

## 1. Introduction

Glutamate decarboxylase (GAD^1^; EC 4.1.1.31) is a tightly regulated cytosolic enzyme in plants that is also widely distributed in animals and bacteria. GAD catalyzes the irreversible decarboxylation of glutamate to γ-aminobutyrate (GABA) and CO_2_. This represents the first committed step of the GABA shunt, a pathway that bypasses the 2-oxoglutarate to succinate reactions of the tricarboxylic acid cycle, and that facilitates plant acclimation to various (a)biotic stresses (Benidickson et al., 2023; Bouché et al., 2003; Che-Othman et al., 2020; Joshi et al., 2019). GADs belong to the aspartate aminotransferase superfamily of pyridoxal-5’-P (PLP)-dependent enzymes (Rossignoli et al., 2018). The genome of the model plant *Arabidopsis thaliana* encodes five closely related GAD isozymes. AtGAD1 (AT5G17330) is the most studied paralog, and is primarily expressed in roots (Benidickson et al., 2023; Bouché et al., 2004; Turano & Fang, 1998). Knockout of AtGAD1 in *atgad1* T-DNA mutants decreases root GABA levels by up to 85% while compromising the ability of Arabidopsis to survive various stresses including salinity, excessive heat, or nutritional Pi deprivation (Benidickson et al., 2023; Mekonnen et al., 2016; Shelp, 1999; Su et al., 2019).

Plant GADs are activated by a reduction in cytosolic pH from resting levels (∼pH 7.3), since their pH-activity optimum is about pH 6.0 (Snedden et al., 1995). However, AtGAD1, AtGAD2, and AtGAD4 each contain a calmodulin (CaM) binding domain near their C-terminus that makes them responsive to cytosolic Ca^2+^, and results in their activation by Ca^2+^/CaM at pH 7.3 (Baum et al., 1993; Shelp et al., 2012; Snedden et al., 1995). The AtGAD1 crystal structure indicates that it occurs as a homohexamer composed of a trimer of dimers, whereas in solution the enzyme’s 58 kDa subunits exist in a dynamic dimer-hexamer equilibrium (Astegno et al., 2015; Gut et al., 2009). The association of dimers into hexamers is promoted by high AtGAD1 concentrations, Ca^2+^/CaM binding, or pH values below 6.5. Tight control of AtGAD1 by Ca^2+^/CaM is critical for ensuring appropriate GABA levels and optimal plant development. Transgenic plants expressing a truncated AtGAD1 mutant lacking its C-terminal CaM binding domain exhibit reduced growth, along with lower glutamate and enhanced GABA levels (Baum et al., 1996; S et al., 2021).

Our recent phosphoproteomic study revealed that AtGAD1 became hyperphosphorylated at several conserved serine residues located near its N-terminus 48 h following resupply of 2 mM Pi to Pi-starved, heterotrophic Arabidopsis suspension cell cultures (Mehta et al., 2021). This highlights a potential link between Pi nutrition and a novel post-translational control mechanism for AtGAD1 via reversible phosphorylation. Preliminary evidence of Raytek (2022) supports the hypothesis that differential phosphorylation inhibits the activity of native AtGAD1 following Pi resupply to Pi-starved Arabidopsis. Phenotypic analyses of *atgad1* T-DNA insertional mutants demonstrated that AtGAD1 and the GABA shunt play important adaptive roles during Arabidopsis acclimation to Pi deprivation (Benidickson et al., 2023).

The present study was initiated to purify recombinant AtGAD1 (rAtGAD1) following its heterologous expression in *Escherichia coli* in order to facilitate further studies of the mechanisms and functions of site-specific AtGAD1 phosphorylation (Mehta et al., 2021; Raytek, 2022). However, the C-terminal CaM-binding domain of rAtGAD1 was determined to be quite susceptible to *in vitro* proteolytic truncation by co-extracted thiol endopeptidase activity. The objective of the current study was to systematically diagnose approaches to overcome this since even very limited proteolysis can trigger dramatic changes to an enzyme’s kinetic and regulatory properties (Plaxton, 2019; Plaxton & Preiss, 1987). Prevention of unwanted proteolysis during enzyme purification usually involves the addition of protease inhibitors or cocktails to homogenization buffers to block proteolysis *in situ*, followed by separation of contaminating proteases from the enzyme of interest by column chromatography (Plaxton, 2019). However, despite testing a wide range of protease inhibitors and cocktails, our results indicate that the most effective way to isolate active, non-proteolyzed rAtGAD1 is to extract and purify the enzyme in the presence of exogenous Ca^2+^/CaM.

## 2. Materials and methods

### 2.1 Heterologous expression of His_6_-rAtGAD1

A culture stab of XL1 blue *Escherichia coli* containing *pET15b*-*AtGAD1* carrying an N-terminal His_6_-tag (Trobacher et al., 2013) was cultured overnight in Luria Bertani (LB) media containing 0.1 mg/ml ampicillin. Plasmid DNA was isolated using the ‘Presto Mini Plasmid Kit’ (Geneaid) according to the manufacturer’s protocol and quantified using a ‘NanoDrop One’ system (Thermo Fischer Scientific). To confirm that the complete *AtGAD1* open reading frame was unmutated and in-frame, Taq polymerase-based PCR with gene-specific forward (5’ GCTCTAGACATATGGTGCTCTCCCAC – 3’) and reverse (5’ – CGGGATCCTTAGCAGATACCACTCG – 3’) primers, and Sanger sequencing (TCAG, SickKids) of *pET15b*-*AtGAD1* was completed. PCR reactions were conducted as follows: initial denaturation at 95 °C for 30 s, followed by 35 cycles of, 95 °C for 20 s, 48 °C for 20 s, and 68 °C for 1 min 45 s, and a final extension at 68 °C for 10 min. Amplicons were visualized on a 1% agarose gel stained with RedSafe Nucleic Acid Staining Solution (FroggaBio). *pET15b*-*AtGAD1* was transformed into *E. coli* BL21 (DE3) CodonPlus-RIL (CP-RIL) or pLysS cells using a standard heat-shock protocol (Green & Sambrook, 2012). Transformants were spread onto LB agar containing 0.1 mg/ml ampicillin and incubated at 37 °C overnight. Individual colonies were selected and cultured overnight at 37 °C, 200 rpm in 5 ml of LB media containing 0.1 mg/ml ampicillin. Plasmids were isolated from individual colonies and confirmed to have the full *AtGAD1* open reading frame, as described above. Transformed cells were cultured in 1.5 L of LB media containing 0.1 mg/ml ampicillin and incubated at 37 °C and 200 rpm until an *A_600_* of 0.5-0.7 was reached. *His_6_-rAtGAD1* expression was induced using 0.1 mM isopropyl β-d-1-thiogalactopyranoside (IPTG) for 4 h at 37 °C and 200 rpm. Cells were harvested by centrifuging at 4,400 × *g* for 15 min, and the resulting pellets frozen in liquid N_2_ and stored at −80 °C.

### 2.2 Purification of His_6_-rAtGAD1

All chromatography steps were performed at 23 °C using an ÄKTA Purifier Fast Protein Liquid Chromatography system (FPLC; Cytiva). Peak FPLC fractions containing purified His_6_-rAtGAD1 were routinely pooled and adjusted to contain 0.1 mM PLP, concentrated with an Amicon Ultra-15 centrifugal filter unit (30 kDa MWCO), desalted (into 50 mM HEPES-KOH, pH 7.2, containing 10% glycerol, 0.5 mM DTT, and 0.1 mM PLP), divided into 50 μL aliquots, frozen in liquid N_2_, and stored at −80 °C.

His_6_-rAtGAD1 was initially purified by following a previously reported procedure (Astegno et al., 2015; Gut et al., 2009; Trobacher et al., 2013). Quick-frozen His_6_-rAtGAD1 expressing *E. coli* BL21 (DE3) CP-RIL cells (7.4 gFW) were thawed and resuspended (1:5; w/v) in buffer A (50 mM HEPES-KOH, pH 7.2, containing 150 mM NaCl, 1 mM DTT, 0.1 mM PLP, 1 mM phenylmethylsulfonyl fluoride [PMSF], and 1X SigmaFAST Protease Inhibitor Cocktail [PIC] Tablets; Millipore-Sigma, cat.# S8830), and lysed by passing twice through a French Pressure cell at 20,000 psi. As PMSF is labile in aqueous solutions (Plaxton, 2019), it was added to extraction buffers immediately prior to cell lysis (from a 100 mM stock prepared in 100% ethanol). Extracts were clarified via centrifugation for 20 min at 48,250 × *g* and 4 °C. The supernatant fluid was adjusted to contain 2 mM CaCl_2_ and batch adsorbed via end-over-end mixing for 30 min at 4 °C with 2 ml of CaM-Sepharose 4B (Cytiva) pre-equilibrated with buffer A containing 2 mM CaCl_2_. The resin was packed into a 1 cm diameter column and connected to the FPLC system. Following elution of nonbinding proteins at 1 ml/min, His_6_-rAtGAD1 was eluted with buffer A containing 2 mM EGTA.

To determine if treatment with 2,2-dipyridyl disulfide (DPDS) suppresses partial rAtGAD1 proteolysis during its extraction and purification, cultures of His_6_-rAtGAD1 expressing *E. coli* BL21 (DE3) CP-RIL cells were prepared as described above. IPTG-induced *pET15b-AtGAD1* BL21 CP-RIL cells (6 gFW) were resuspended (1:5; w/v) in buffer B (20 mM NaH_2_PO_4_, pH 7.4, 300 mM NaCl, 10 mM imidazole, 0.1 mM PLP) containing 2 mM DPDS, lysed using the French press and centrifuged as described above. The supernatant fluid was loaded at 1 ml/min onto a column (1 × 5 cm) of HisPur Ni-NTA Superflow Agarose (Thermo Fisher Scientific) pre-equilibrated with buffer B lacking 0.1 mM PLP. After eluting non-binding proteins with buffer C (20 mM NaH_2_PO_4_, pH 7.4, 300 mM NaCl, and 20 mM imidazole), His_6_-rAtGAD1 was eluted using buffer C containing 300 mM imidazole.

To determine whether Ca^2+^/CaM addition suppresses *in vitro* proteolysis of His_6_-rAtGAD1, cultures of *E. coli* BL21 (DE3) pLysS cells expressing petunia CaM81 (exhibits 100% sequence identity to Arabidopsis CaM7) in the *pET5a* expression vector (Fromm & N.-H., 1992; Teresinski et al., 2023) were prepared as described above. IPTG-induced *pET15b-AtGAD1* BL21 CP-RIL and *pET5a-CaM81* BL21 pLysS cells (6 gFW each) were combined and resuspended (1:5; w/v) in buffer B containing 1 mM CaCl_2_, lysed using the French press and centrifuged as described above. The supernatant fluid was loaded at 1 ml/min onto a column (1 × 5 cm) of HisPur Ni-NTA Superflow Agarose pre-equilibrated with buffer B containing 0.1 mM CaCl_2_ and lacking 0.1 mM PLP. After eluting non-binding proteins with buffer C containing 0.1 mM CaCl_2_, His_6_-rAtGAD1 and CaM81 were eluted using the same buffer containing 300 mM imidazole.

### 2.3 Protein concentration determination, electrophoresis, and immunoblotting

Protein concentrations were determined using the Pierce BCA Protein Assay Kit (Thermo Fischer Scientific) with a bovine γ-globulin standard. SDS-PAGE was performed using a Bio-Rad Mini-PROTEAN 3 Cell mini-gel rig, and stacking and separating gels composed of 4 and 10% (w/v) acrylamide, respectively, unless otherwise indicated. Following electrophoresis, gels were either stained for total protein with Coomassie Brilliant Blue G-250 or R-250, or electroblotted onto a PVDF membrane for immunoblotting. Immunoblots were probed for 1 h at 23 °C with rabbit immune serum raised against petunia GAD (anti-GAD) (Arazi et al., 1995) or mouse anti-recombinant *Petunia* × *hybrida* CaM72 monoclonal IgG (anti-CaM) (Baum et al., 1996), or overnight at 4 °C with mouse anti-His-IgG (anti-His, Cell Signaling). Immunoreactive polypeptides were visualized using an alkaline phosphatase-linked secondary antibody and chromogenic detection. For the determination of His_6_-rAtGAD1 subunit molecular mass during SDS-PAGE a plot of Relative Mobility versus log *M_r_* was constructed using Bio-Rad’s Precision Plus Protein™ Standards (Cat. # 1610394) having the following *M_r_* values: 150, 100, 75, 50, 37 and 25 kDa.

### 2.4 GAD activity assays

GAD activity was assayed using a coupled GABase spectrophotometric assay as previously described (Miyashita & Good, 2008). GABase was obtained from MilliporeSigma (Product #: G7509) and consists of a mixture of GABA transaminase and succinic semialdehyde dehydrogenase from *Pseudomonas fluorescens*. Unless otherwise stated, the stopped-time assay was carried out at 25 °C for 30 min in a 200 μL reaction mixture containing 25 mM MES and 25 mM bis-tris propane (pH 5.8 or 7.3), 10% (v/v) glycerol, 10 mM KCl, 20 mM glutamate, 1 mM DTT and 0.5 mM PLP. Assays were terminated by bringing the assay to pH 1.0 with 70% perchloric acid, followed by adjustment to pH 8.6 with 3 M KOH. Samples were centrifuged for 3 min at 17,500 × *g*, from which up to 100 μL of supernatant fluid was transferred to a 96-well microtiter plate. A 100 μL aliquot of a GABase reaction mixture (100 mM bis-tris-propane, pH 8.6, 10 mM 2-oxoglutarate, 2 mM NADP^+^, 0.16 units/ml GABase) was added to each well. A Molecular Devices Spectromax Plus Microplate spectrophotometer was used to monitor the increase in *A_340_* owing to the reduction of NADP^+^ to NADPH. Using a calibration curve generated with various amounts of GABA (Supplemental Fig. S1A), the specific activity of His_6_-rAtGAD1 was calculated as units/mg protein, where one unit is defined as 1 μmol of GABA produced per min at 25 °C. The assays were validated by showing that the amount of GABA produced was proportional to assay time (Supplemental Fig. S1B) and GAD concentration.

### 2.5 Statistical analysis

Data was analyzed using two-way ANOVA and Tukey’s multiple comparison tests. Results were deemed statistically significant at *p* < 0.05.

## 3. Results and discussion

### 3.1 The C-terminal region of His_6_-rAtGAD1 is susceptible to partial in vitro proteolytic truncation

His_6_-rAtGAD1 was heterologously expressed in *E. coli* BL21 (DE3) CP-RIL and extracted under denaturing conditions in hot SDS-PAGE sample buffer. Anti-GAD or anti-His immunoblotting following SDS-PAGE of the resulting extract revealed a single immunoreactive 59 kDa polypeptide (Fig. 1A and B), which is consistent with the enzyme’s predicted subunit *M_r_* (i.e., AtGAD1 = 502 amino acids, plus N-terminal His_6_ tag and linker = 21 amino acids). His_6_-rAtGAD1 was then extracted and purified under non-denaturing conditions by following a well established protocol (Astegno et al., 2015; Gut et al., 2009; Trobacher et al., 2013). This includes adding 1 mM PMSF and SigmaFAST PIC to the extraction buffer^2^, and subjecting the resulting soluble extract to CaM-Sepharose affinity FPLC. This purification yielded 1.8 mg of purified His_6_-rAtGAD1 having a specific activity of 15.3 ±1.1 units/mg (mean ±SEM of *n* = 3 determinations) at optimal pH 5.8. However, SDS-PAGE and immunoblotting of the final preparation revealed an approximate 1:1 ratio of anti-GAD and anti-His immunoreactive, and protein-staining polypeptides of 59 and 54 kDa (p59 and p54, respectively) (Fig. 1A-C). This indicates that partial degradation of the p59 to p54 occurred *in vitro* following His_6_-rAtGAD1 extraction. The anti-His immunoblots also demonstrate that the N-terminal His_6_-tag of purified, partially degraded His_6_-rAtGAD1 subunits (i.e. the p54) remained intact (Fig. 1B, lane 2). Thus, *in vitro* truncation of p59 to p54 (Fig. 1, lane 2) arises from cleavage of a susceptible peptide bond within the enzyme’s C-terminal region, which encompasses its CaM binding domain (Fig. 2). As the partially degraded His_6_-rAtGAD1 effectively bound to CaM-Sepharose 4B in the presence of Ca^2+^, this indicates that at least two rAtGAD1 subunits (i.e. the p59) in the multimeric protein remained intact. AtGAD1’s crystal structure revealed that a highly flexible alpha-helical linker region (residues 449–470) connects its autoinhibitory CaM binding domain (residues 471–502) to the enzyme’s core (Gut et al., 2009). The partial proteolysis that we observed likely occurs near the middle of this flexible linker region; i.e., cleavage of His_6_-rAtGAD1’s p59 subunits around Gln459 (Fig. 2) would yield the p54. It is notable that a similar protein staining ‘doublet’ was reported following SDS-PAGE of purified His_6_-rAtGAD1 that had been expressed in *E. coli* or Sf9 insect cells (Trobacher et al., 2013; Zik et al., 1998). By contrast, evidence of proteolytically truncated His_6_-rAtGAD1 was not presented in several reports concerning its expression in *E. coli* BL21 (DE3) pLysS and purification (Astegno et al., 2015; Gut et al., 2009; Turano & Fang, 1998). Nevertheless, potential ‘protease issues’ were alluded to by Turano and Fang (1998) who stated that “*Crude protein extracts were assayed at pH 5.8 and not 7.3 because there were problems associated with CaM binding, and the effects of proteases on the CaM-binding domain in these extracts*”. Similarly, Trobacher et al. (2013) mentioned that: “*Assays of GAD activity and determination of spectral properties were simultaneously conducted as soon as possible after the recombinant protein was extracted to minimize proteolysis that seemed to occur even in the presence of several popular protease inhibitor*s”.

**Fig. 1.**
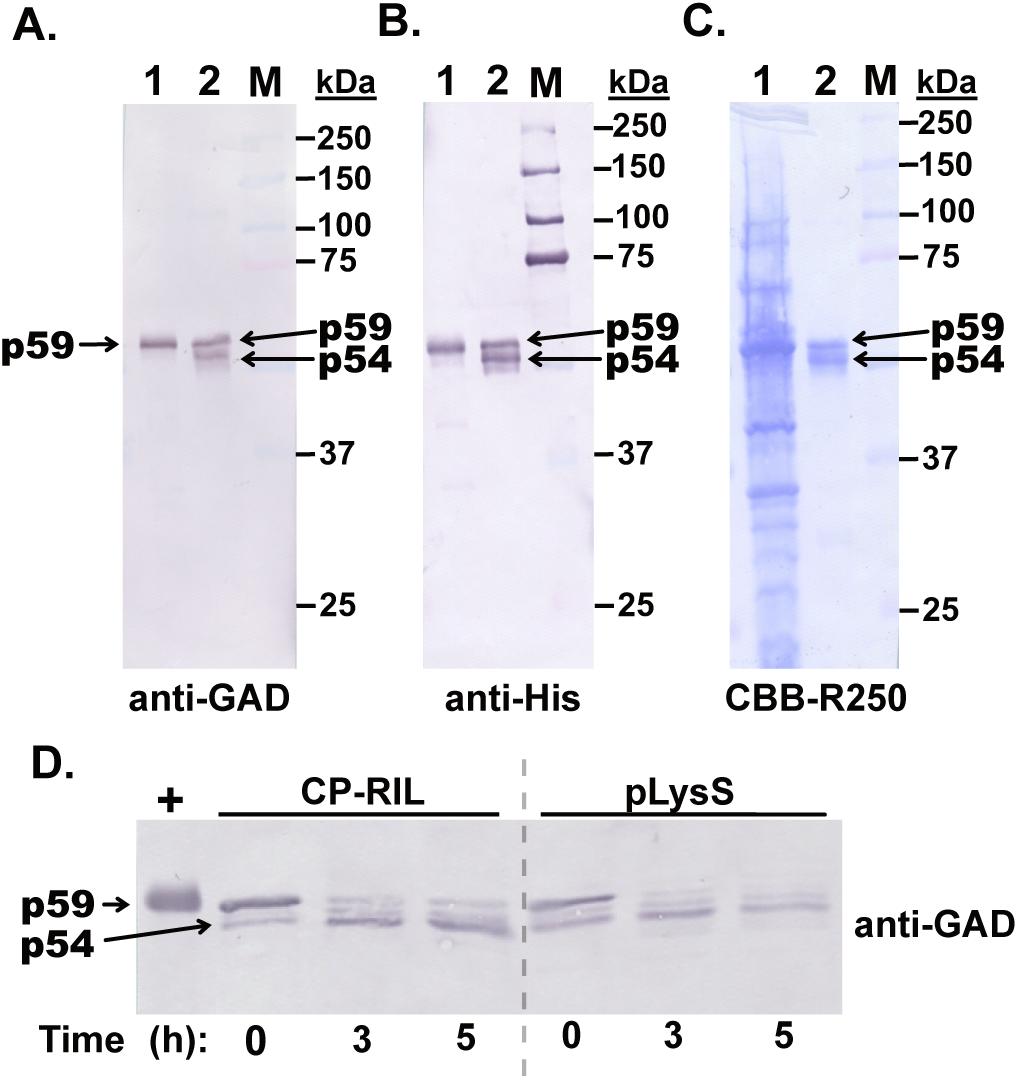
rAtGAD1 is susceptible to *in vitro* proteolytic truncation following its heterologous expression in *E. coli*. SDS-PAGE was followed by anti-GAD (A, D) or anti-His (B) immunoblotting, or (C) total protein staining with Coomassie Brilliant Blue R-250 (CBB-R250). Lane 1 of panels A-C contains 2 µL of a 10-fold diluted (A, B) or undiluted (C) *pET15b-AtGAD1* BL21 CP-RIL extract prepared under denaturing conditions in hot SDS-PAGE sample buffer. Lane 2 of panels A-C contains 50 ng (A, B) or 1 µg (C) of His_6_-rAtGAD1 extracted under non-denaturing conditions from *pET15b-AtGAD1* BL21 CP-RIL cells in presence of 1 mM PMSF and 1X SigmaFAST PIC, and purified via CaM-Sepharose affinity chromatography as previously described (Astegno et al., 2015; Gut et al., 2009; Trobacher et al., 2013). ‘M’ denotes various *M_r_*standards. (D) Clarified extracts were prepared from *pET15b-AtGAD1* BL21 CP-RIL or pLysS cells in 50 mM HEPES-KOH (pH 7.2) containing 150 mM NaCl, 1 mM DTT, 0.1 mM PLP, 1 mM PMSF and 1X SigmaFAST PIC, and incubated at 23 °C. Aliquots were removed at the indicated times and subjected to SDS-PAGE and immunoblotting using anti-GAD (10 µg protein loaded/lane). The ‘+’ lane of panel D contains 2 µL of a 10-fold diluted *pET15b-AtGAD1* BL21 CP-RIL extract prepared under denaturing conditions in hot SDS-PAGE sample buffer.

**Fig. 2.**
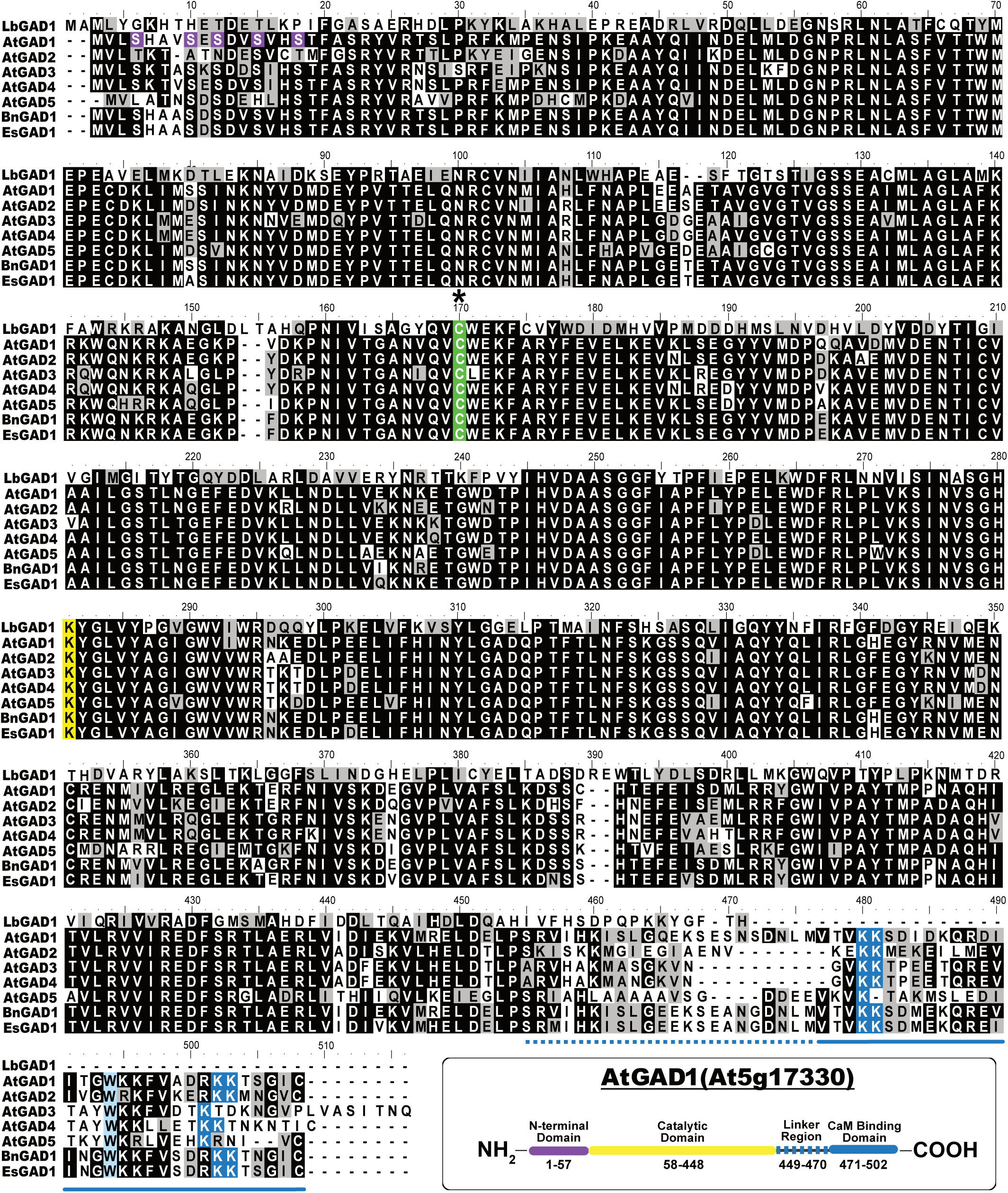
Amino acid sequence alignment of AtGAD1 with its paralogs and several orthologs, highlighting residues and domains critical to GAD function. The PLP-binding Lys277 residue of AtGAD1 is highlighted in yellow, whereas its conserved Cys166 residue that aligns with the PLP-anchoring Cys168 of *Lactobacillus* GAD (Huang et al., 2018; Sun et al., 2021) is highlighted in green and marked by an asterisk. The regions corresponding to AtGAD1’s C-terminal CaM-binding domain and its adjacent flexible ‘linker region’ (Gut *et al.,* 2009) are underlined with solid and dotted lines, respectively. A key tryptophan residue of the CaM-binding domain is highlighted in light blue, and flanking CaM-anchoring lysine residues (Snedden et al., 1995) highlighted in dark blue. AtGAD1’s N-terminal phosphoserine residues identified during LC-MS/MS analysis of Pi-resupplied *Arabidopsis* cell cultures (Mehta *et al*., 2021) are highlighted in purple. The deduced GAD sequences were aligned using ClustalW (http://www.ebi.ac.uk/Tools/clustalw2/index.html). Identical and similar amino acids are indicated by black and grey shading, respectively. Inset: schematic diagram of AtGAD1 functional domains (Gut et al., 2009).

To further assess the apparent inability of PMSF or the SigmaFAST PIC to suppress partial *in vitro* proteolysis of His_6_-rAtGAD1 (Fig. 1A-C), clarified extracts from His_6_-rAtGAD1-expressing *E. coli* BL21 (DE3) CP-RIL or pLysS cells were prepared in their presence and incubated at 23 °C. Aliquots were removed at 0, 3 and 5 h, boiled in SDS-PAGE sample buffer, and subjected to immunoblotting with anti-GAD. Initially, a prominent immunoreactive p59 corresponding to non-degraded His_6_-rAtGAD was observed (Fig. 1D). However, incubation at 23 °C led to the progressive disappearance of the p59 and concomitant appearance of the p54, similar to that observed for the purified, partially proteolyzed His_6_-rAtGAD1 preparation. The combined results of Fig. 1 demonstrate that His_6_-rAtGAD1 is vulnerable to partial proteolysis of its C-terminal, CaM-binding domain by co-extracted protease activity that: (i) is insensitive to either PMSF, or any of the protease inhibitors present in SigmaFAST PIC tablets, and (ii) occurs irrespective of the particular His_6_-rAtGAD1expressing *E. coli* BL21 (DE3) strain being employed (i.e. CP-RIL or pLysS).

### 3.2 The thiol modifiers 2,2’-dipiridyl disulfide and N-ethylmaleimide block in vitro His_6_-rAtGAD1 proteolysis, but also abolish its GAD activity

Various class-specific protease inhibitors and commercially available PICs (Supplemental Table S1) were tested for their ability to prevent *in vitro* proteolysis of His_6_-rAtGAD1 during incubation of *pET15b-AtGAD1* BL21 (DE3) CP-RIL *E. coli* extracts at 23 °C. As occurred with PMSF and the SigmaFAST PIC (Fig. 1), the following substances were ineffective as judged by anti-GAD immunoblotting of extracts incubated for up to 24 h: 1 mM *p*-hydroxymecuribenzoate, 1 mM iodoacetamide, 10 µM E-64, 100 µM tosyl phenylalanyl chloromethyl ketone, 100 µM tosyl lysyl chloromethyl ketone, 5 mM 1,10-phenanthroline, 5 mM EDTA, 10 µg/mL leupeptin, 10 µg/mL chymostatin, 10 µg/mL pepstatin A, 5 µL/mL ProteCEASE-100 PIC (G-Biosciences), or 1x Roche Complete PIC tablet (MilliporeSigma) (Supplemental Fig. S2). In contrast, *in vitro* p59 proteolysis was blocked when *E. coli* extraction was performed in the presence of the thiol modifiers DPDS (2 mM) or N-ethylmaleimide (NEM) (10 mM) (Fig. 3A; Supplemental Fig. S2). Unlike NEM which is a non-specific thiol modifier (Plaxton, 2019), DPDS was reported to specifically react with active site thiol groups of native, but not denatured, papain or peptidase A (well characterized cysteine proteases from papaya fruit) ((Baines & Brocklehurst, 1982; Brocklehurst & Little, 1973). We therefore extracted His_6_-rAtGAD1 with 2 mM DPDS, and purified 1.4 mg of the enzyme via Ni^2+^-affinity FPLC. SDS-PAGE and anti-GAD immunoblotting of the final preparation indicated that the His_6_-rAtGAD1 was purified to near homogeneity, and that DPDS conferred long-term protection of the enzyme’s p59 subunits to partial *in vitro* proteolysis (Fig. 3B and 3C). However, spectrophotometric GAD activity assays unexpectedly revealed that the purified, non-proteolyzed His_6_-rAtGAD1 that had been extracted in presence of 2 mM DPDS was catalytically inactive. We also assessed the impact of 2 mM DPDS or 10 mM NEM inclusion in the extraction buffer on GAD activity in the resulting clarified extracts. DPDS or NEM addition caused a complete loss of GAD activity, as opposed to control extracts prepared in the absence of either reagent that exhibited a GAD specific activity of 0.45 ±0.01 units/mg (mean ±SEM of *n* = 3 determinations) at pH 5.8.

**Fig. 3.**
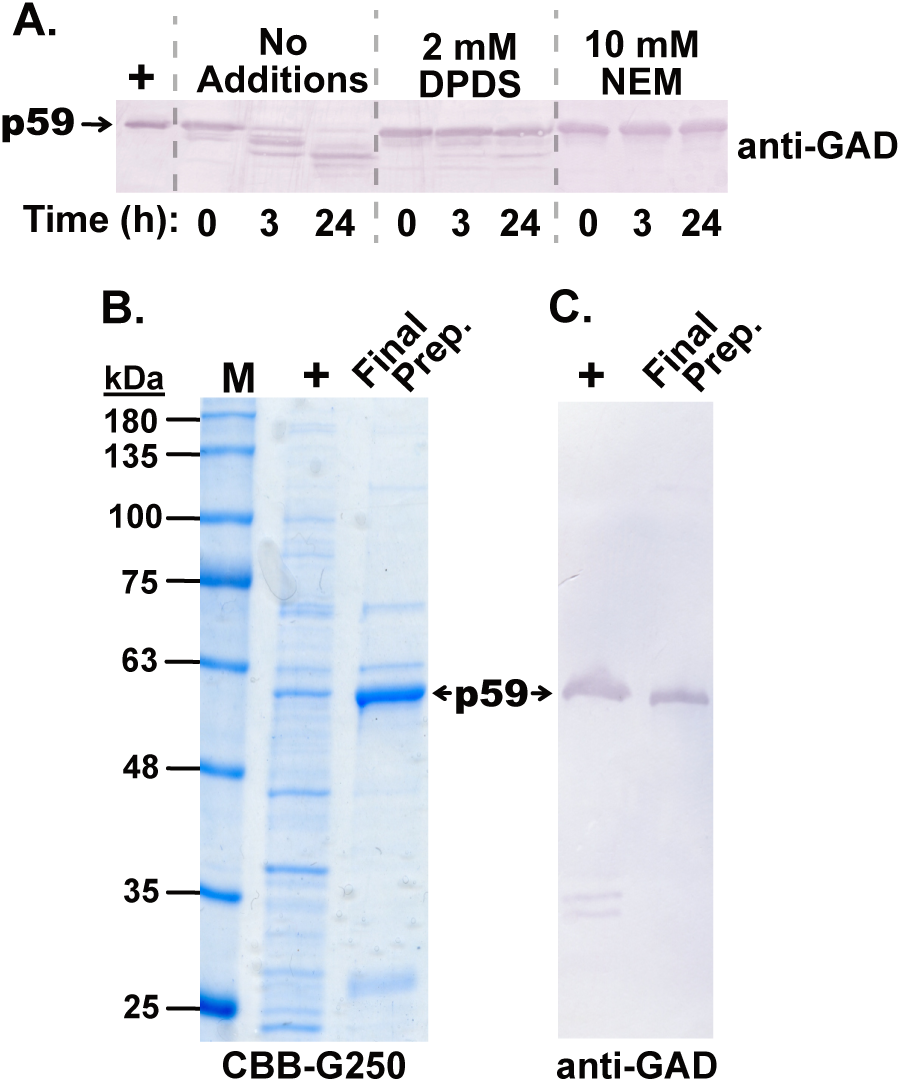
The thiol modifiers DPDS and NEM prevent *in vitro* proteolysis of His_6_-rAtGAD1. (A) Clarified extracts from *pET15b-AtGAD1* BL21 CP-RIL cells were incubated at 23 °C in the absence and presence of 2 mM DPDS or 10 mM NEM. Aliquots were removed at the indicated times and subjected to SDS-PAGE and immunoblotting using anti-GAD (10 µg protein loaded/lane). (B, C) SDS-PAGE and immunoblot analysis of purified, non-proteolyzed His_6_-rAtGAD1. His_6_-rAtGAD1 was extracted from *pET15b-AtGAD1* BL21 CP-RIL cells in presence of 2 mM DPDS and purified via Ni^2+^-affinity FPLC as described in the Materials and Methods. SDS-PAGE of 1 µg and 50 ng of the final preparation was respectively followed by (B) protein staining with Coomassie Brilliant Blue G-250 (CBB-G250) and (C) immunoblotting with anti-GAD. The ‘+’ lanes of panels A-C contain 2 µL of a 10-fold diluted *pET15b-AtGAD1* BL21 CP-RIL extract prepared under denaturing conditions in hot SDS-PAGE sample buffer, whereas ‘M’ denotes various *M_r_* standards.

These results indicate that DPDS and NEM modified a key sulfhydryl group of His_6_-rAtGAD1 that is essential for catalytic activity, resulting in inactivation. This is supported by the observation that thiol-directed reagents inactivate a purified potato GAD (Satyanarayan & Nair, 1985). Furthermore, structural studies revealed that Cys168 participates in anchoring the PLP cofactor to the active site of *Lactobacillus brevis* GAD, thereby supporting catalytic activity (Huang et al., 2018). Sequence alignment indicated that Cys166 may be the site of rAtGAD1 modification in the presence of DPDS or NEM, since it aligns with Cys168 of the *Lactobacillus* GAD, while being conserved in AtGAD1’s paralogs and orthologs (Fig. 2). Although DPDS-inactivated papain or peptidase A could be reactivated following treatment with 50 mM β-mercaptoethanol at pH 8.0 (Baines & Brocklehurst, 1982; Brocklehurst & Little, 1973), we were unable to recover any GAD activity following incubation of the purified, DPDS-treated His_6_-rAtGAD1 with 50 mM β-mercaptoethanol or 10 mM DTT for up to 30 min at pH 8.0 and 23 °C.

### 3.3. Ca^2+^/CaM addition blocks in vitro proteolytic truncation of His_6_-rAtGAD1

As the C-terminal CaM-binding domain of His_6_-rAtGAD1 is prone to truncation by co-extracted *E. coli* protease activity (Fig. 1), and ligand addition can protect certain enzymes from *in vitro* proteolysis (Plaxton, 2019), we assessed whether Ca**^2+^**/CaM addition to the extraction buffer might be an effective strategy to obtain non-proteolyzed, active His_6_-rAtGAD1. Structural studies show that Ca^2+^/CaM activates AtGAD1 in a distinctive way by relieving two C-terminal autoinhibitory domains of adjacent active sites, forming a 393 kDa AtGAD1–CaM heterohexameric complex with an unusual 1:3 CaM:AtGAD1 stoichiometry (Gut et al., 2009).

Anti-GAD immunoblotting demonstrated that *in vitro* p59 proteolysis was largely nullified when *pET15b-AtGAD1* BL21 CP-RIL *E. coli* cells were co-extracted with an equivalent amount of *pET5a-CaM81* BL21 pLysS cells and 1 mM CaCl_2_, and the resulting clarified extract incubated at 23 °C for up to 5 h (Fig. 4A). SDS-PAGE and anti-GAD immunoblotting of His_6_-rAtGAD1 extracted and purified via Ni^2+^-affinity FPLC in the presence of Ca^2+^/CaM81 revealed a single prominent protein-staining and immunoreactive p59 (Fig. 4B and 4C). This indicates that the His_6_-rAtGAD1 was purified to near homogeneity, and that its co-extraction and purification in the presence of CaM81-expressing *E. coli* cells and CaCl_2_ is an efficient approach to impede its proteolytic truncation. This purification yielded 2.3 mg of non-proteolyzed His_6_-rAtGAD1 having a specific activity of 26.6 ±1.1 units/mg (mean ±SEM of *n* = 3 determinations) at pH 5.8 (Fig. 5). Anti-CaM immunoblotting confirmed the presence of a 15 kDa immunoreactive CaM81 polypeptide (lacks His-tag) in the Ni^2+^-affinity FPLC-purified His_6_-rAtGAD1 preparation (Fig. 4D).

**Fig. 4.**
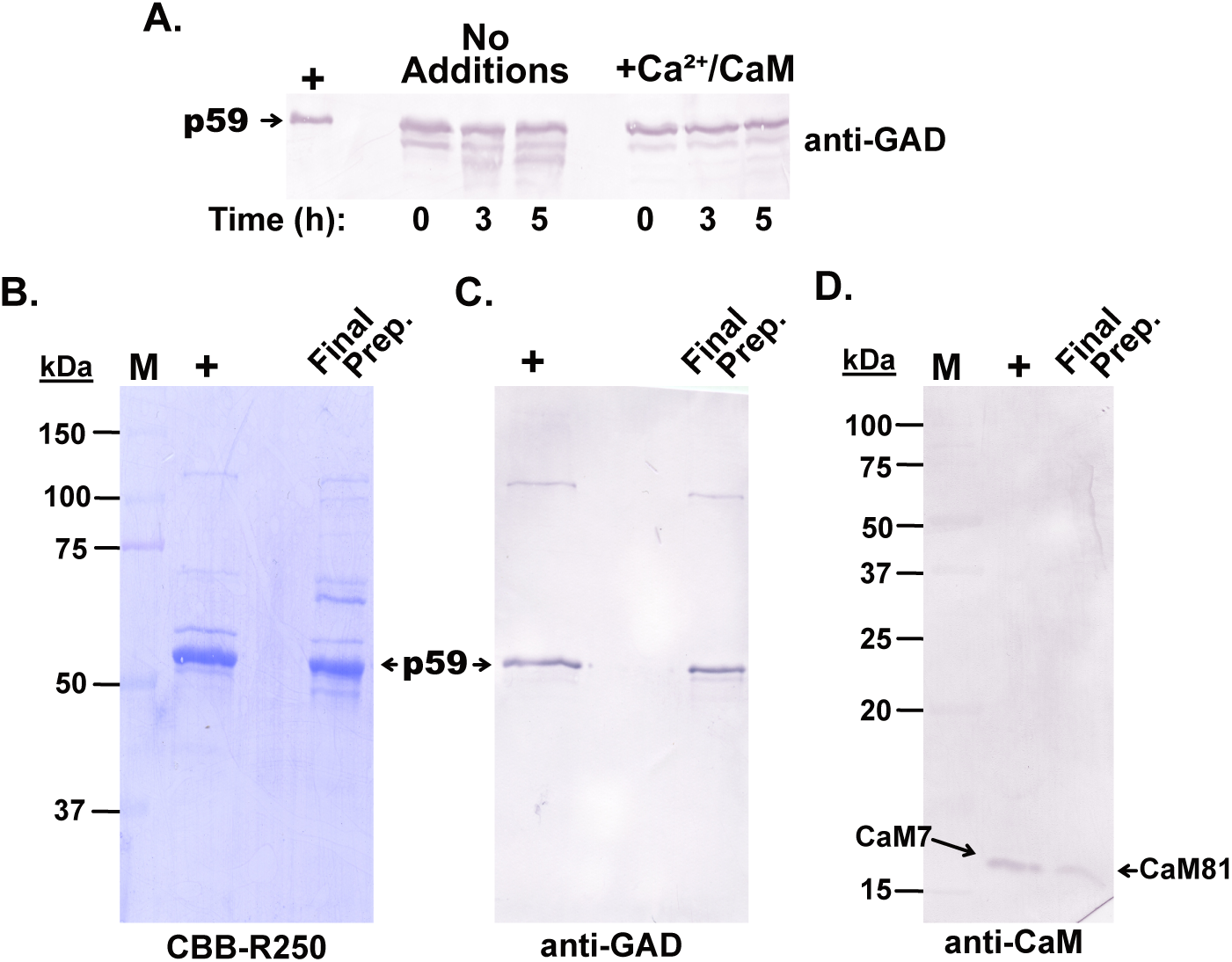
Ca^2+^/CaM addition inhibits proteolytic truncation of His_6_-rAtGAD1. Equivalent amounts of *pET15b-AtGAD1* BL21 CP-RIL and *pET5a-CaM81* BL21 pLysS cells were co-extracted in the presence of 1 mM CaCl_2_. (A) The clarified extract was incubated at 23 ℃, and aliquots removed at the indicated times and subjected to SDS-PAGE and immunoblotting using anti-GAD (10 µg protein loaded/lane). (B-D) The His_6_-rAtGAD1:CaM81 complex was purified via Ni^2+^-affinity FPLC and analyzed by SDS-PAGE followed by: (B) Coomassie Brilliant Blue R-250 (CBB R-250) protein staining (1 µg/lane), and immunoblotting with (C) anti-GAD (50 ng/lane) or (D) anti-CaM (200 ng/lane). The ‘**+**’ lanes of panels A-C denote non-proteolyzed His_6_-rAtGAD1 (100 ng for panel A, 1 µg for panel B, 50 ng for panel C) extracted in the presence of 2 mM DPDS and purified via Ni^2+^-affinity FPLC, whereas the ‘**+**’ lane of panel D was purified recombinant Arabidopsis CaM7 (50 ng) (Vanderbeld & Snedden, 2007). ‘M’ of panels B and D denotes various *M_r_* standards. The SDS-PAGE resolving gel acrylamide concentration was increased from 10 to 15% for the anti-CaM immunoblot shown in panel D.

**Fig. 5.**
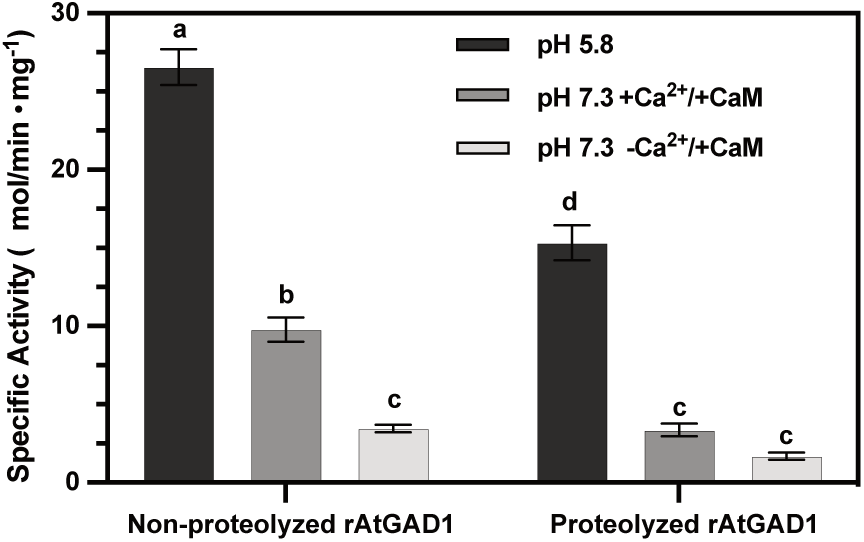
Partial proteolysis reduces the activity and Ca^2+^/CaM sensitivity of His_6_-rAtGAD1. Spectrophotometric (GABase coupled) activity assays were conducted as outlined in the Materials and Methods with 130 and 260 ng of non-proteolyzed and partially proteolyzed rAtGAD1, respectively, at pH 5.8 and pH 7.3. Ca^2+^/CaM-sensitivity was tested at pH 7.3 by the addition 1 µM petunia CaM81 in the presence of either 1 mM CaCl_2_ (+Ca^2+^/+CaM) or 5 mM EGTA (-Ca^2+^/+CaM). Proteolyzed and non-proteolyzed His_6_-rAtGAD1 respectively correspond to the final preparations shown in Fig. 1A-C and Fig. 4B and C. All values represent mean ±SEM of *n* = 3 separate assays; different letters indicate significant differences (*p* < 0.05; two-way ANOVA) between the proteolyzed and non-proteolyzed rAtGAD1 preparations.

### 3.4. Partial proteolysis reduces the activity and Ca^2+^/CaM sensitivity of purified His_6_-rAtGAD1

Activities of purified partially proteolyzed and non-proteolyzed His_6_-rAtGAD1 were compared at optimal pH 5.8, and at pH 7.3 in the presence of 0.1 µM petunia CaM81 ±1 mM CaCl_2_ (Fig. 5). Partial *in vitro* proteolysis of p59 to p54 His_6_-rAtGAD1 subunits (Fig. 1) significantly reduced the enzyme’s specific activity at both pH values by at least 40%. Furthermore, although both preparations were activated by Ca^2+^/CaM at pH 7.3, this was significantly more pronounced for non-proteolyzed relative to the partially proteolyzed His_6_-rAtGAD1 which were respectively activated 283 ±19% and 202 ±14% (means ±SEM of *n* = 3 determinations) by Ca^2+^/CaM, relative to control assays containing CaM that lacked Ca^2+^ (Fig. 5).

## 4. Concluding remarks

The detection and prevention of proteolysis during native or recombinant enzyme purification is a concern for all biochemists, particularly when the enzyme of interest has been only partially degraded and retains catalytic activity. Limited proteolysis can result in erroneous conclusions regarding an enzyme’s kinetic and regulatory properties, as documented in the current manuscript (Fig. 5) and numerous prior studies (Plaxton, 2019). The *E. coli* BL21 (DE3) strain employed in the present study is widely used for recombinant protein expression owing to its deficiency in the cytoplasmic serine protease *lon*, and the outer membrane aspartyl protease *ompT*, which may help reduce proteolysis of the target protein (Francis & Page, 2010). However, a co-extracted *E. coli* cysteine endopeptidase appears to be responsible for the *in vitro* truncation of an approximate 5 kDa polypeptide from the C-terminus of His_6_-rAtGAD1’s p59 subunits, since this was prevented when extraction was performed in the presence of the thiol modifiers NEM or DPDS (Fig. 3). NEM irreversibly and non-specifically alkylates protein thiol groups, including active site thiol groups of cysteine proteases such as papain and papain-like proteases (Clementz et al., 2010; Nakajima et al., 1991; Saha et al., 2018). Inactivation of His_6_-rAtGAD1 by NEM was not surprising since prior studies have established the requirement of a key thiol group for plant and microbial GAD catalysis (Huang et al., 2018; Satyanarayan & Nair, 1985). However, His_6_-rAtGAD1 inactivation by DPDS was quite unexpected since unlike NEM, DPDS was reported to function as a substrate analogue that specifically results in covalent (thiopyridone) modification of active site thiol groups of native, but not denatured, papain or peptidase A from papaya fruit (Baines & Brocklehurst, 1982; Brocklehurst & Little, 1973). DPDS has since been successfully employed to block partial proteolysis of active allosteric regulatory enzymes from various plant and green algal sources (e.g., phosphoenolpyruvate carboxylase, phosphoenolpyruvate carboxykinase, pyruvate kinase) by co-extracted cysteine endopeptidase activity (Martín et al., 2007; Plaxton, 2019). However, the current results indicate that caution should be exerted using DPDS as a routine cysteine protease inhibitor since it can apparently not only modify thiol groups at the active site of ‘papain-like’ cysteine proteases, but can also modify thiol groups of the target enzyme being investigated. As DPDS simultaneously inactivated His_6_-rAtGAD1 while protecting the enzyme from proteolysis (Fig. 3), it will be interesting to assess the intriguing possibility that p59 to p54 conversion arises from AtGAD1 autoproteolytic activity, as opposed to a co-extracted *E. coli* cysteine endopeptidase. Autoproteolysis is a known regulatory mechanism for certain enzymes, allowing them to switch between active and inactive states or to modulate their activity in response to specific cellular signals or environmental conditions.

Extraction and Ni^2+^-affinity FPLC in the presence of Ca^2+^/CaM81 proved to be an unconventional, yet effective strategy for the isolation of non-proteolyzed and catalytically active His_6_-rAtGAD1 (Figs. 4 and 5). These results are expected to facilitate studies of the mechanisms and functions of multisite AtGAD1phosphorylation (Mehta et al., 2021; Raytek, 2022). Purified His_6_-rAtGAD1 could be employed as a substrate for assaying and characterizing the protein kinase that phosphorylates AtGAD1 following Pi resupply to Pi-starved Arabidopsis suspension cells (Mehta et al., 2019). Moreover, comparing the kinetic and regulatory properties of non-proteolyzed wild-type His_6_-rAtGAD1 with the corresponding site-directed phosphomimetic mutant could provide invaluable information on the functions of *in vivo* multisite AtGAD1 phosphorylation. This may, in turn, expedite biotechnological approaches to bioengineer Pi-efficient crops, promoting agricultural sustainability and ecosystem preservation, while contributing to future food security.

## Footnotes

1 *Abbreviations*: CaM, calmodulin; CP-RIL; CodonPlus-RIL; DPDS, 2,2′-dipyridyl disulfide; DTT, dithiothreitol; FPLC, fast protein liquid chromatography; GABA, γ-aminobutyrate; GAD, glutamate decarboxylase; IPTG, isopropyl β-d-1-thiogalactopyranoside; LB, Luria Bertani; NEM, N-ethylmaleimide; p54 and p59, 55 and 59 kDa polypeptides, respectively; PIC, protease inhibitor cocktail; PLP, pyridoxal-5’-phosphate; PMSF, phenylmethylsulfonyl fluoride; rAtGAD1, recombinant Arabidopsis GAD1.

2 PMSF irreversibly inactivates serine and certain cysteine proteases (Plaxton, 2019), whereas the SigmaFAST PIC is intended to inhibit serine, cysteine, aspartic and metalloproteases as it contains 4-(2-aminoethyl)benzenesulfonyl fluoride hydrochloride, bestatin, L-(trans)-epoxysuccinyl-leucylamido (4-guanidino) butane (E-64), pepstatin A, phosphoramidon, leupeptin, and aprotinin.

