## Supplemental Table S1 for "Heterologous expression and purification of glutamate decarboxylase-1 from the model plant *Arabidopsis thaliana*: characterization of the enzyme’s *in vitro* truncation by thiol endopeptidase activity"

Protease inhibitors and cocktails analyzed in time course-based diagnosis of partial *in vitro* proteolysis of $His_6$-rAtGAD1. Protease inhibitor stock solutions and protease inhibitor cocktails (PICs) were prepared as described by Plaxton (2019) and according to the PIC manufacturer's recommended protocol, respectively.

| Protease inhibitor[a] or cocktail | Protease specificity | Mode of action | Final concentration |
|---|---|---|---|
| EDTA | Metallo- | Reversible | 5 mM |
| 1,10-Phenanthroline | Metallo- | Reversible | 5 mM |
| Pepstatin A | Aspartic | Reversible | 10 μg/ml |
| Chymostatin | Serine & cysteine | Reversible | 10 μg/ml |
| Leupeptin | Serine & cysteine | Reversible | 10 μg/ml |
| PMSF | Serine & cysteine | Reversible | 2 mM |
| TLCK | Serine & cysteine | Irreversible | 100 μM |
| TPCK | Serine & cysteine | Irreversible | 100 μM |
| DPDS | Cysteine | Irreversible | 2 mM |
| E-64 | Cysteine | Irreversible | 10 μM |
| pHMB | Cysteine | Irreversible | 1 mM |
| Iodoacetamide | Cysteine | Reversible | 1 mM |
| NEM | Cysteine | Irreversible | 10 mM |
| ProteCEASE-100 PIC (G-Biosciences) | Cysteine, serine & metallo- | Reversible & irreversible | 1x |
| Roche cOmplete PIC tablets (Millipore-Sigma) | Cysteine, serine & metallo- | Reversible & irreversible | 1x |
| SIGMAFAST PIC tablets (Millipore-Sigma) | Cysteine, serine, aspartic, & metallo- | Reversible & irreversible | 1x |

[a]Abbreviation: DPDS, 2,2'-dipyridyl disulfide; E-64, L-trans-expoxysuccinyl-leucylamide-(4-guanido)-butane; EDTA, ethylenediaminetetraacetic acid; pHMB, *p*-hydroxymercuribenzoate; NEM, N-ethylmaleimide; PMSF, phenylmethylsulfonyl fluoride; TLCK, tosyl lysyl chloromethyl ketone; TPCK, tosyl phenylalanyl chloromethyl ketone.

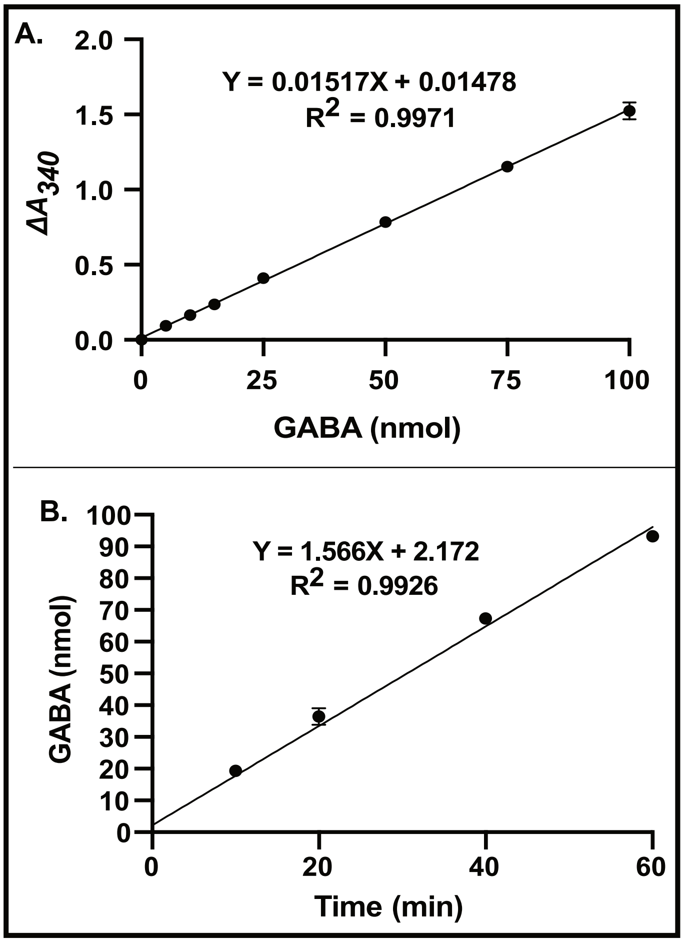


**Supplemental Fig. S1. Calibration and validation of the spectrophotometric GABase-linked GAD activity assay.** (A) GABase calibration curve generated with the specified amounts of GABA. (B) The GABase-coupled GAD assay was conducted as described in the Materials and Methods and validated by demonstrating that the amount of GABA produced by 130 ng of purified $His_6$-rAtGAD1 was proportional to reaction time. All values in panels A and B represent means (±SEM) of $n$ = 3 separate assays, with linear regression values indicated for each plot; where invisible the SEMs are too small to be seen.

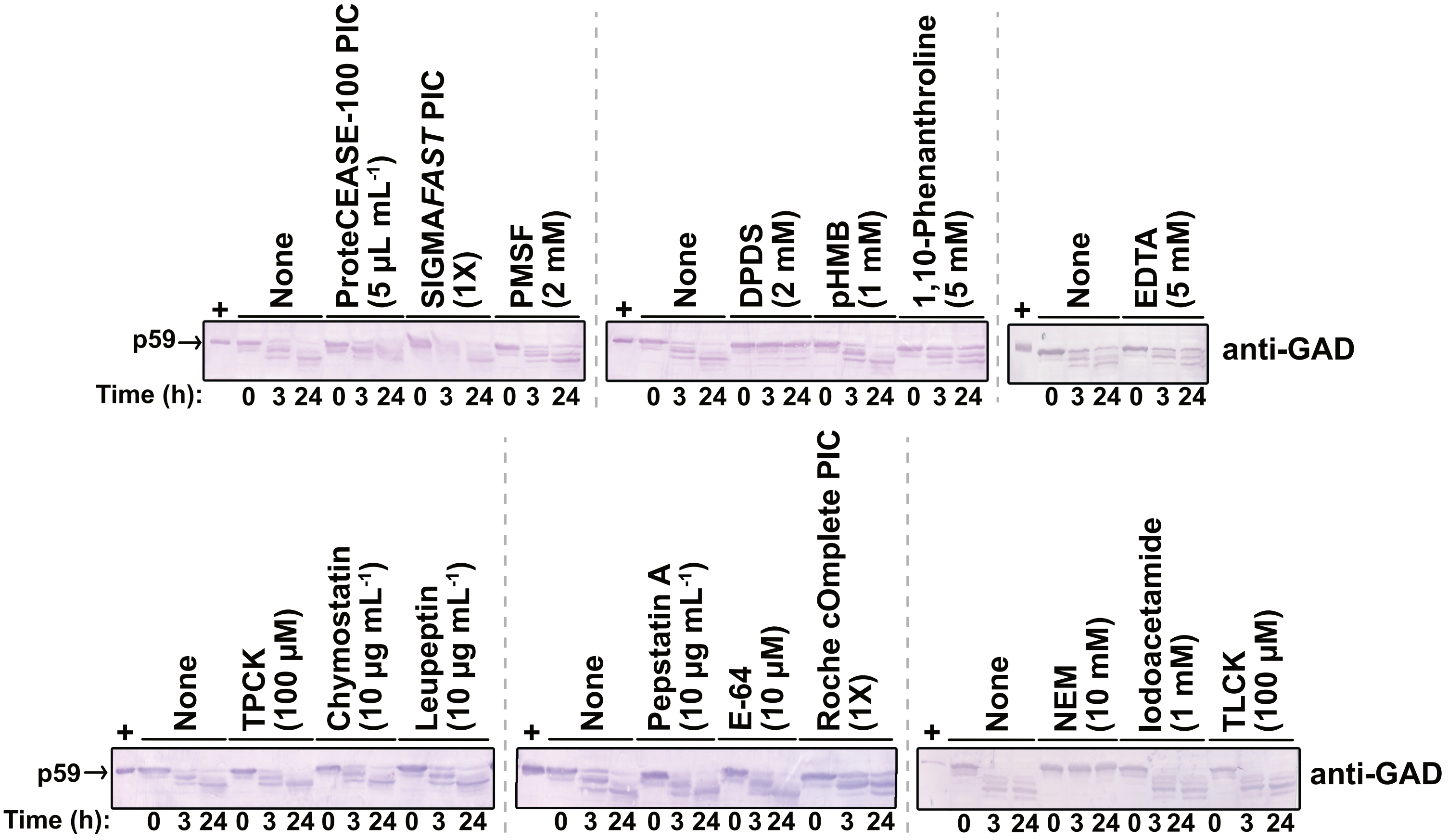


**Supplemental Fig. S2. Influence of various protease inhibitors and cocktails on *in vitro* proteolytic susceptibility of the $His_6$-rAtGAD1 p59.** Clarified extracts were prepared under non-denaturing conditions from *pET15b-AtGAD1* BL21 (CP-RIL) *E. coli* cells in the presence and absence of the protease inhibitors and cocktails listed in Supplemental Table S1. Aliquots were removed at the indicated times and subjected to SDS-PAGE and immunoblotting using anti-GAD (10 µg protein/lane). The '**+**' lanes denote 2 µL of a 10-fold diluted extract prepared in hot SDS sample buffer.
